# Additive and interactive effects of spatial attention and expectation on perceptual decisions

**DOI:** 10.1101/235218

**Authors:** Arianna Zuanazzi, Uta Noppeney

**Author notes:** Correspondence: Arianna Zuanazzi, Computational Cognitive Neuroimaging Laboratory, Computational Neuroscience and Cognitive Robotics Centre, University of Birmingham, B15 2TT Birmingham, UK.

## Abstract

Spatial attention and expectation are two critical top-down mechanisms controlling perceptual inference. Based on previous research it remains unclear whether their influence on perceptual decisions is additive or interactive.

We developed a novel multisensory approach that orthogonally manipulated spatial attention (i.e. task relevance) and expectation (i.e. signal probability) selectively in audition and evaluated their effects on observers’ responses in vision. Critically, while experiment 1 manipulated expectation directly via the probability of task-relevant auditory targets across hemifields, experiment 2 manipulated it indirectly via task-irrelevant auditory non-targets.

Surprisingly, our results demonstrate that spatial attention and signal probability influence perceptual decisions either additively or interactively. These seemingly contradictory results can be explained parsimoniously by a model that combines spatial attention, general and spatially selective response probabilities as predictors with no direct influence of signal probability. Our model provides a novel perspective on how spatial attention and expectations facilitate effective interactions with the environment.

---

Generating a coherent representation of the world from the sensory signals with which we are bombarded is a fundamental challenge facing us in everyday life. Crucially, perceptual inference is not purely driven by bottom-up signals but also guided by top-down selective attention and prior expectations (or predictions). Selective attention shapes perception by prioritizing processing of signals that are relevant for our current goals. Conversely, prior expectations encode the probabilistic structure of the environment^1^. Based on past events and experiences we generate expectations or predictions for the future. While attention and expectation are both thought to facilitate perception at the behavioural level^2-10^, they have traditionally been associated with distinct neural effects. Attention amplifies neural activity and signal to noise ratio^11-14^ while expectation often leads to a reduction in stimulus-induced neural activity^15-20^.

According to the notion of predictive coding, perceptual inference emerges within the cortical hierarchy via iterative adjustment of top-down predictions against bottom-up sensory evidence. While backwards connections impose predictions from higher to subordinate level, forwards connections furnish the prediction error, i.e. the discrepancy between prediction and sensory evidence, from lower to higher hierarchical levels^21-25^. Attention may influence perceptual inference by enhancing the precision (i.e. inverse of variance) of the prediction and/or prediction error signal leading to an increase in sensory gain for attended signals^24-26^. As a consequence, attention and expectation jointly shape perceptual inference and decisions.

Surprisingly, research to date has mostly conflated attention and expectation^8,27,28^. Most prominently, the so-called Posner cuing paradigm^29^ manipulates observers’ endogenous spatial attention using a cue that probabilistically predicts the location of the subsequent signal thereby confounding spatial attention and spatial expectation (i.e. signal probability)^30^. Only recently have studies attempted to dissociate spatial attention and expectation by orthogonally manipulating task-relevance (i.e. response requirement) and spatial signal probability. Using fMRI, a previous study by Kok et al.^31^ showed that spatial attention and expectation influence neural responses in an interactive fashion. More specifically, attention reversed the activation increase for unexpected relative to expected signals that were observed for unattended signals. The interactive effects between attention and spatial signal probability were interpreted as in line with the notion of precision weighted prediction errors as embodied in predictive coding models^24,31^. Yet, a critical limitation of those neuroimaging experiments is that synergistic effects between attention and expectation could be evaluated only at the neural but not the behavioural level^31,32^, because spatial attention was manipulated as response requirement over space^32^. As a result, observers did not respond to the unattended, i.e. task-irrelevant, signals and the effects of spatial expectation on response times could only be evaluated for signals in the attended hemifield^31^. This raises the critical question whether the interactions between spatial attention and spatial signal probability are behaviourally relevant for effective interactions with the environment. How does the brain optimize detection of signals across the spatial field depending on attention and expectations formed based on signal probability? In the current study we have developed a novel multisensory approach to determine whether spatial attention and signal probability influence behaviour additively and/or interactively in a target detection task.

## Results

In a series of two experiments, participants were presented on each trial with an auditory burst of white noise or a visual flash in their left or right hemifields. We orthogonally manipulated spatial attention (i.e. response requirement for signals presented in a particular hemifield) and expectation (i.e. signal probability in a particular hemifield) selectively in audition and evaluated their effects on target detection in audition and vision, where signal probability and response requirements were held constant. This multisensory generalization approach^27,28^ provides us with the novel opportunity to evaluate the putative additive or interactive effects of spatial attention and signal probability at the behavioural level rather than only implicitly at the neural level as in previous unisensory research^31^.

Critically, experiment 1 manipulated auditory spatial expectation directly via the probability of auditory targets across the two hemifields, which led to differences in the general response probability across conditions (i.e. run type A (blue) vs run type B (green), see Figure 1a, 1b and 1d, Supplementary Table S1 and ^31^ for related design). By contrast, experiment 2 manipulated auditory expectation via task-irrelevant non-targets that never required a response and was thereby able to hold the general response probability constant across all conditions (see Figure 2a, 2b and 2d, Supplementary Table S1 and ^32^ for related design).

**Fig. 1.**
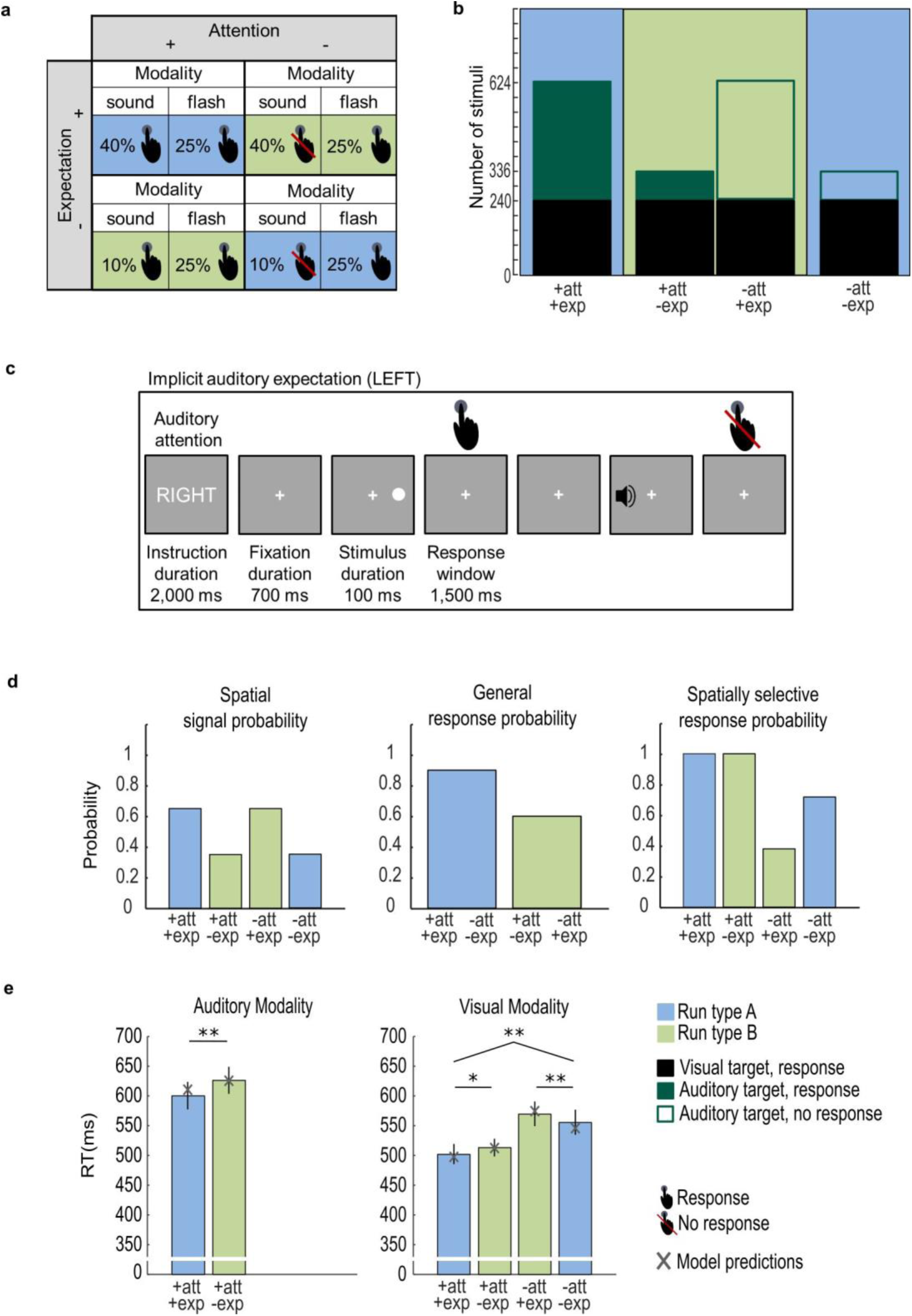
Experiment 1: Design, example trial, probabilities, observed and predicted response times. **a.** Auditory spatial attention and expectation (i.e. signal probability) were manipulated in a 2 (attended vs. unattended) × 2 (expected vs. unexpected) × 2 (auditory vs. visual) factorial design. Colours indicate run type (blue: attention and expectation are congruent; green: attention and expectation are incongruent). Presence vs. absence of response requirement is indicated by the hand symbol. **b**. Number of auditory (dark green) and visual (black) trials in the 2 (attended vs. unattended) × 2 (expected vs. unexpected) conditions. The bar plots are filled (i.e. response requirement) or not filled (i.e. no response requirement). The fraction of filled area for each bar represents the spatially selective response probability. The fraction of filled area pooled over the two bars of one particular run type (e.g. blue) represents the general response probability in a run; it is greater for the ‘blue runs’ where attention and expectation are congruent. **c.** At the beginning of each run, a cue informed participants whether to attend and respond to auditory signals selectively in their left or right hemifield throughout the entire run. On each trial participants were presented with an auditory or visual stimulus (100 ms duration) either in their left or right hemifield. They were instructed to respond to auditory stimuli only in the ‘attended’ hemifield and to all visual stimuli irrespective of hemifield as fast and accurately as possible with the same finger. The response window was limited to 1500 ms. Participants were not explicitly informed that auditory signals were more likely in one of the two hemifields. Instead, spatial expectation (i.e. spatial signal probability) was implicitly learnt over runs. **d.** The bar plots show i. spatial signal probability: the probability that a signal (pooled over visual and auditory modalities) was presented in a particular hemifield, ii. general response probability: the probability that a trial required a response in a particular (green or blue) run type, iii. spatially selective response probability: the probability that a signal required a response conditioned on that it was presented in a particular hemifield. **e.** The bar plots show across subjects’ mean (±SEM) response times for each of the six conditions with response requirements. The brackets and stars indicate significance of main effects and interactions. * *p* < 0.05; ** *p* < 0.01. The grey crosses show the condition-specific response times predicted by the ‘winning’ Model_Log_ 2 that includes attention, general response probability and spatially selective response probability as predictors.

**Fig. 2.**
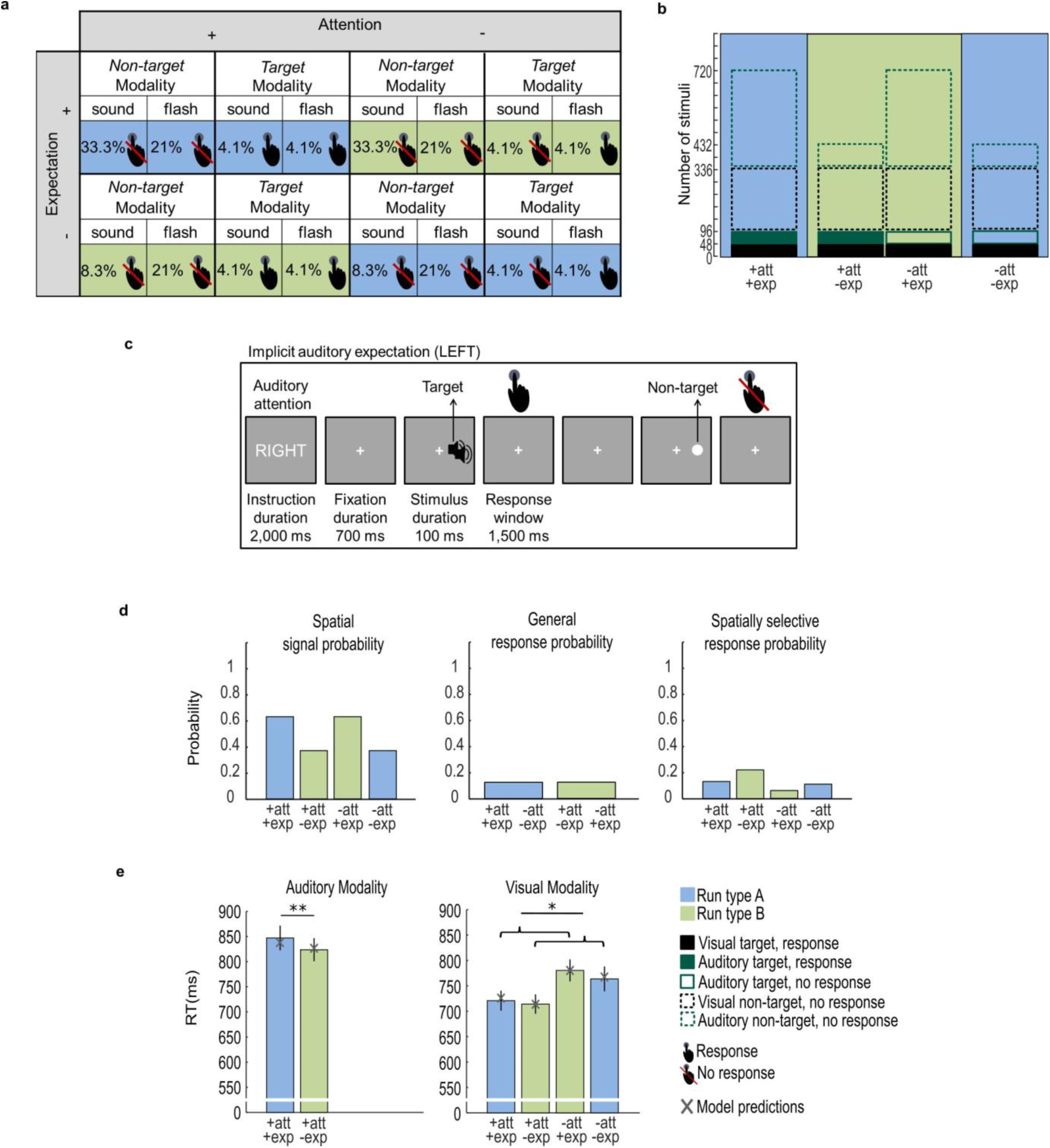
Experiment 2: Design, example trial, probabilities, observed and predicted response times. **a.** Auditory spatial attention and expectation (i.e. signal probability) were manipulated in a 2 (attended vs. unattended) × 2 (expected vs. unexpected) × 2 (auditory vs. visual) × 2 (target vs. non-target) factorial design. Colours indicate run type (blue: attention and expectation are congruent; green: attention and expectation are incongruent). Presence vs. absence of response requirement is indicated by the hand symbol. **b**. Number of auditory (dark green) and visual (black) trials in the 2 (attended vs. unattended) × 2 (expected vs. unexpected) conditions. The bar plots are filled (i.e. response required) or not filled (i.e. no response required). Their contours indicate target vs. non-target trials (solid = target; dotted = non-target). The fraction of filled area for each bar represents the spatially selective response probability, which is greater in the unexpected than expected conditions. **c.** At the beginning of each run, a cue informed participants whether to attend and respond to auditory targets (i.e. double noise burst) selectively in their left or right hemifield throughout the entire run. On each trial participants were presented with an auditory or visual stimulus (100 ms duration) either in their left or right hemifield. They were instructed to respond to auditory targets (i.e. double noise bursts) only in the ‘attended’ hemifield and to all visual targets (i.e. double flash) irrespective of hemifield as fast and accurately as possible with the same finger. The visual or auditory non-targets never required any response. The response window was limited to 1500 ms. Participants were not explicitly informed that auditory signals were more likely in one of the two hemifields. Instead, spatial expectation (i.e. spatial signal probability) was implicitly learnt over runs. **d.** The bar plots show i. spatial signal probability: the probability that a signal (pooled over visual and auditory modalities) was presented in a particular hemifield, ii. general response probability: the probability that a trial required a response in a particular (green or blue) run type, iii. spatially selective response probability: the probability that a signal required a response conditioned on that it was presented in a particular hemifield. **e.** The bar plot shows across subjects’ mean (±SEM) response times for each of the six conditions with response requirements. The brackets and stars indicate significance of main effects and interactions. * *p* < 0.05; ** *p* < 0.01. The grey crosses show the condition-specific response times predicted by the ‘winning’ Model_Log_ 2 that includes attention, general response probability and spatially selective response probability as predictors.

First, we asked for each experiment independently whether spatial attention and signal probability shapes target detection responses in an additive or interactive fashion. This question can be addressed only for visual signals where responses were collected over both attended and unattended hemifields. To assess whether the effects of expectation generalise from audition to vision in the attended hemifield, we also report the results for auditory targets in the attended hemifield.

### Experiment 1

For the auditory modality, the two-sided paired-sample t-tests on hit rates and response times in the attended hemifield showed non-significantly higher hit rates (*t*(14) = 2.06, *p* = 0.058, Cohen’s *d*_*av*_ [95% CI] = 0.56 [0.02, 1.12]) and significantly faster responses when the hemifield was expected than unexpected (*t*(14) = −3.23, *p*=.006, Cohen’s *d*_*av*_ [95% CI] = −0.31 [−0.52, −0.09] (see Supplementary Figure S1, left bar plot of Figure 1e and Table 1).

**Table 1.** Group mean reaction times (RT) and hit rates for each stimulus modality in each condition for experiment 1 and experiment 2. D-prime and bias for the visual modality in each condition for experiment 2. Please note that d-prime and bias could be computed neither for experiment 1 (because there were no non-targets included in the paradigm) nor for auditory modality of experiment 2 (because participants did not make any false alarms in the +att-exp condition).

|  | Auditory modality |  | Visual modality |  |  |  |
| --- | --- | --- | --- | --- | --- | --- |
|  | +att<br>+exp | +att<br>-exp | +att<br>+exp | +att<br>-exp | -att<br>+exp | -att<br>-exp |
| Experiment 1 |  |  |  |  |  |  |
| RT (ms)<br>(SEM) | 599.9<br>(22.8) | 626.8<br>(22.8) | 502.5<br>(16.1) | 514.2<br>(14.4) | 567.3<br>(20.6) | 553.2<br>(21) |
| Hit rates (%)<br>(SEM) | 99.4<br>(0.2) | 98.4<br>(0.6) | 99.5<br>(0.2) | 98.9<br>(0.4) | 98.8<br>(0.4) | 99.2<br>(0.3) |
| Experiment 2 |  |  |  |  |  |  |
| RT (ms)<br>(SEM) | 847.2<br>(24.6) | 823.8<br>(23.0) | 720.8<br>(19.8) | 714.2<br>(19.2) | 780.4<br>(21.7) | 763.7<br>(24.5) |
| Hit rates (%)<br>(SEM) | 97.6<br>(0.8) | 98.3<br>(0.6) | 96<br>(0.9) | 97<br>(0.8) | 96.2<br>(0.9) | 96.4<br>(0.9) |
| D-prime<br>(SEM) | / | / | 4.08<br>(0.12) | 4.07<br>(0.15) | 4.25<br>(0.12) | 4.12<br>(0.15) |
| Bias<br>(SEM) | / | / | 0.19<br>(0.07) | 0.08<br>(0.05) | 0.24<br>(0.06) | 0.17<br>(0.05) |

Note. Standard errors of the mean (SEM) are given in parentheses.

For the visual modality, the 2 (attended vs. unattended) × 2 (expected vs. unexpected) repeated measures ANOVAs revealed a significant main effect of attention on hit rates (*F*(1, 14) = 5.56, *p* = 0.033, 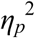 [90% CI] = 0.29 [0.01, 0.51]) and response times (*F*(1, 14) = 42.81, *p* < 0.001, 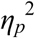 [90% CI] = 0.75 [0.48, 0.84]). Critically, we also observed a significant crossover interaction between attention and expectation for hit rates (*F*(1, 14) = 4.89, *p* = 0.044, 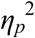 [90% CI] = 0.26 [0.004, 0.49]) and response times (*F*(1, 14) = 8.90, *p* = 0.010, 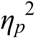 [90% CI] = 0.39 [0.06, 0.59]). Participants had non-significantly higher hit rates (*t*(14) = 2.01, *p* = 0.064, Cohen’s *d*_*av*_ [95% CI] = 0.44 [-0.02, 0.90]) and responded significantly faster (*t*(14) = −2.16, *p* = 0.049, Cohen’s *d*_*av*_ [95% CI] = −0.2 [−0.39, −0.001]) to visual targets in the attended hemifield when this hemifield was expected than unexpected (see Supplementary Figure S1, right bar plot of Figure 1e and Table 1). This simple main effect mimics the response profile observed for auditory targets, suggesting that the effect of expectation generalises from audition to vision. By contrast, participants had non-significantly lower hit rates (*t*(14) = −1.49, *p* = 0.159, Cohen’s *d*_*av*_ [95% CI] = −0.23 [−0.53, 0.09]) and responded significantly more slowly (*t*(14) = 3.21, *p* = 0.006, Cohen’s *d*_*av*_ [95% CI] = 0.18 [0.05, 0.30]) to visual targets in the unattended hemifield, when this hemifield was expected than unexpected (see Supplementary Figure S1, right bar plot of Figure 1e and Table 1).

### Experiment 2

For the auditory modality, the two-sided paired-sample t-tests showed slower responses (*t*(20) = 3.43, *p* = 0.003, Cohen’s *d*_*av*_ [95% CI] = 0.21 [0.07, 0.35]) in the attended hemifield, when this hemifield was expected than unexpected (see left bar plot of Figure 2e and Table 1).

For the visual modality, the 2 (attended vs. unattended) × 2 (expected vs. unexpected) repeated measures ANOVA on response times revealed a significant main effect of attention (*F*(1, 20) = 62.58, *p* < 0.001, 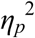 [90% CI] = 0.76 [0.55, 0.83]). As in experiment 1, participants responded faster to visual stimuli in their attended than unattended hemifield. Yet, in contrast to experiment 1, we did not observe a significant interaction between attention and expectation in experiment 2. Instead, the repeated measures ANOVA revealed a significant main effect of expectation for both bias (*F*(1, 20) = 5.17, *p* = 0.034, 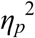 [90% CI] = 0.20 [0.009, 0.42]) and response times (*F*(1, 20) = 5.23, *p* = 0.033, 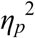 [90% CI] = 0.21 [0.009, 0.42]). Irrespective of whether the hemifield was attended or unattended, participants had a greater bias and responded more slowly to visual stimuli in the expected than unexpected spatial hemifield, again mimicking the effect of expectation in the auditory modality (see Supplementary Figure S1, right bar plot of Figure 2e and Table 1).

In summary, even though both experimental paradigms orthogonally manipulated spatial attention and expectation operationally defined as signal probability, they led us to strikingly different conclusions. Experiment 1 suggests that attention and expectation act in an interactive fashion: under spatial attention, participants responded faster to auditory and visual targets in their expected hemifield, but this effect was reversed for visual targets when attention is diverted to the other hemifield. By contrast, experiment 2 suggests that spatial attention and expectation influence response times in an additive fashion. Here, participants were generally slower when responding to targets in the expected than unexpected hemifield irrespective of attention. Not only does this main effect of expectation in experiment 2 contradict the findings from experiment 1, but it may also be surprising given the vast literature suggesting that expectation facilitates processing^3,8,33^. These contradictory results across the two experiments suggest that modelling the experiments as a 2 (attended vs. unattended) × 2 (expected vs. unexpected) factorial design may not provide a coherent explanatory framework.

Key to resolving these seemingly contradictory results is the realization that spatial attention and signal probability jointly determine two additional probabilities critical for perceptual decision making: 1. the general response probability, i.e. the probability that a response is required on a given trial irrespective of where the stimulus is presented and 2. the spatially selective response probability, i.e. the probability that a response is required conditioned on a stimulus being presented in a particular hemifield (see Figure 1d and 2d and Supplementary Table S1). In the following we therefore investigate whether models that include general and spatially selective response probability outperform the traditional factorial model of attention and expectation in accounting for observers’ response times jointly across both experiments.

### Joint models of experiment 1 and experiment 2

We generated a 3 (combination of fixed effects predictors) × 2 (linear vs. log-transform of probability values in the predictors) model space (see methods section for details and Figure 3). All models assume that the observer prioritises processing in hemifields that are explicitly attended via task instructions per se, yet they differ in whether expectation (i.e. spatial signal probability), general response probability or spatially selective response probability are included as additional explanatory variables. Model 1 is the conventional factorial model that allows for additive and interactive effects of attention and signal probability. Model 2 assumes that signal probability per se does not directly affect response times, but only indirectly by co-determining the general and spatially selective response probability. Model 3 is based on Model 2, but it is more complex by allowing signal probability to influence response times not only indirectly, by co-determining general and spatially selective response probability, but also directly. In addition, given previous research showing a non-linear relationship between probabilities and response times^34^, we manipulated whether probabilities predict response times in a linear (i.e. Model_Lin_) or log-transformed (i.e. Model_Log_) fashion.

**Fig. 3.**
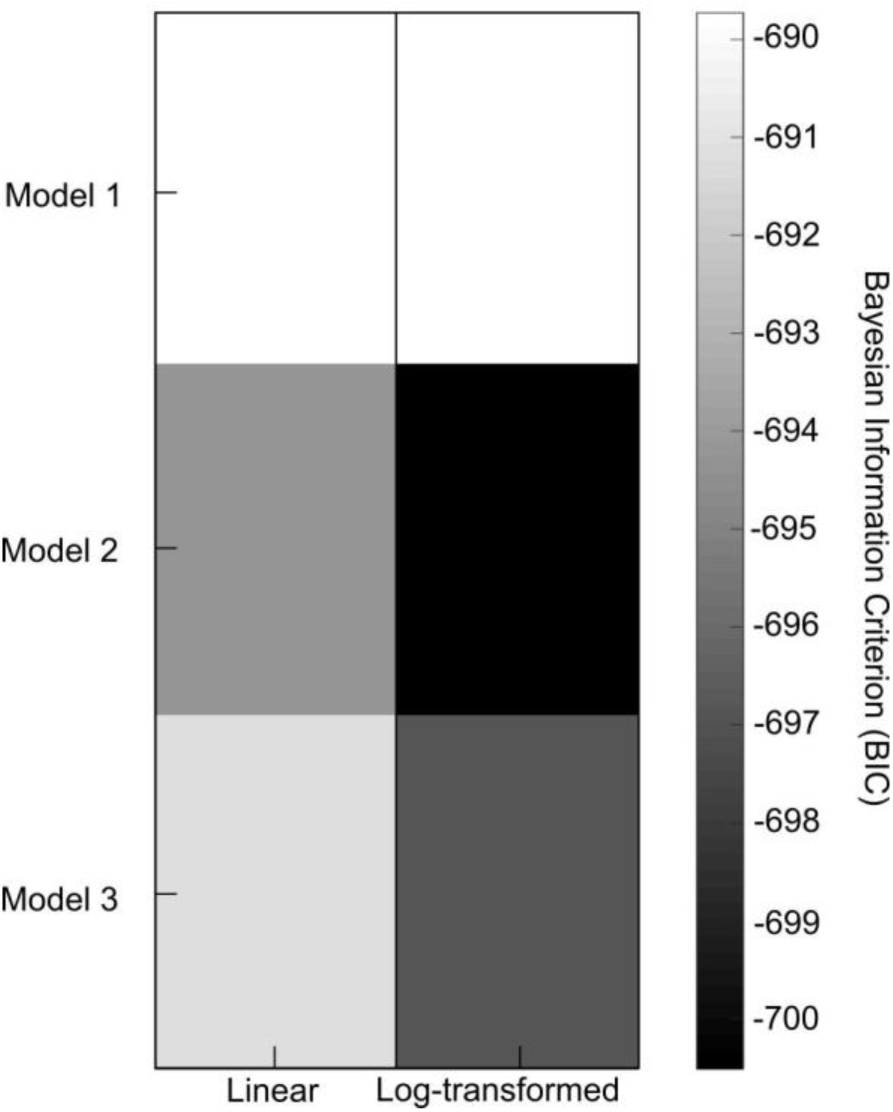
Factorial model space and Bayesian Information Criterion (BIC) for the joint analysis of experiment 1 and experiment 2. 3 (combination of fixed effects predictors) × 2 (probability values in predictors: linear vs. log-transformed) model space. Predictors included in i. conventional Model 1: attention, spatial signal probability and attention × spatial signal probability interaction, ii. Model 2: attention, general response probability and spatially selective response probability, iii. Model 3: attention, spatial signal probability, general response probability and spatially selective response probability. To account for potential non-linear relationship between probabilities and response times, the probabilities predicted response times in a linear or a log-transformed fashion. The matrix shows the Bayesian Information Criterion (BIC) of the 6 models (smaller BIC, i.e. darker shade, indicates better model).

Figure 3 shows the Bayesian Information Criterion values for the six models in a factorial matrix (Bayesian Information criterion (BIC): Model_Lin_ 1: −690; Model_Lin_ 2: −694; Model_Lin_ 3: −691; Model_Log_ 1: −690; Model_Log_ 2: −701; Model_Log_ 3: −697; n.b. a smaller value indicates a better model). First, we show that log-transformed probabilities are better predictors for response times in our target detection task than probabilities per se^34^. Second and more importantly, Model_Log_ 2 that combines spatial attention with general and spatially selective response probability outperformed both the conventional factorial model as well as a more elaborate model that also includes spatial signal probability as a predictor by itself (see Figure 3). More specifically, Bayesian model comparison provided strong evidence (i.e. increase in BIC of approx. 10) for the winning Model_Log_ 2 relative to the conventional Model_Log_ 1 and Model_Log_ 3. Figures 1e and 2e show the response time predictions of the winning Model_Log_ 2 as grey crosses together with observed condition-specific response times. In contrast to the conventional Model 1, Model_Log_ 2 can flexibly model that response times in the attended hemifield are faster for expected than unexpected stimuli in experiment 1, but slower in experiment 2. Critically, adding spatial signal probability in Model 3 did not increase the model evidence. These results suggest that expectation (i.e. spatial signal probability) influences response times mainly indirectly by co-determining general and spatially selective response probabilities.

One may argue that the spatially selective response probability already accounts for observers’ allocation of attention over space. However, a seventh model, i.e. the winning Model_Log_ 2 without the attention predictor, obtained a BIC of only −686 suggesting that endogenous attention influences perceptual decisions as an independent predictor above and beyond spatially selective response probability.

## Discussion

Prior expectations and attention are two key determinants controlling perceptual inference. The current study developed a novel multisensory approach to dissociate additive and interactive effects of attention and expectation at the behavioural level. Manipulating attention and signal probability selectively in the auditory modality, we evaluated their effects on signal detection in the auditory and visual modalities. In each experiment, we observed qualitatively equivalent effects in audition and vision for signal probability in the attended hemifield demonstrating the effectiveness of multisensory generalization in our experiments^27,28^. Critically however, even though both experiments manipulated spatial attention and signal probability orthogonally, they revealed either interactive (experiment 1) or additive (experiment 2) effects.

In experiment 1, we observed synergistic interactions between spatial attention and signal probability at the behavioural level consistent with previously reported interactive effects at the neural level in an equivalent experimental paradigm^31^. When stimuli were attended, participants responded faster to signals presented in the expected than unexpected hemifield, which is in line with the general notion that prior expectations facilitate perceptual processing^35^. Yet, when signals were unattended, stimuli presented in the expected hemifield were associated with slower response times than in the unexpected hemifield. This crossover interaction between spatial attention and signal probability can be explained by the effects of general response probability (and spatially selective response probability). As shown in figure 1d (middle bar plot) and in Supplementary Table S1, in runs (blue) where attention and expectation are directed to the same spatial hemifields, participants are required to respond on 90% of the trials as compared to 60% of the trials in runs when attention and expectation are directed to different hemifields.

Experiment 2 manipulated spatial signal probability via auditory non-targets that never required any response. As a consequence, the general response probability was constant across conditions (see Figure 2d middle bar plot, Supplementary Table S1 and ^32^ for related design). Further, increasing spatial signal probability via an increase in the number of non-targets for expected relative to unexpected hemifields (see Figure 2a and 2b) necessarily decreases the spatially selective response probability (Figure 2d, right bar plot). Consistent with the profile of the spatially selective response probability, but contrary to conventional views of expectation^36^, signals presented in the expected relative to unexpected hemifield were generally associated with slower response times. Put simply, the neural system does not facilitate target detection in the spatial hemifield where many events occur (i.e. high signal probability, Figure 2d left bar plot), but in the hemifield where a high percentage of those events requires a response (i.e. a high spatially selective response probability, Figure 2d, right bar plot). In line with this qualitative explanation, formal Bayesian model comparison confirmed that the response time profiles across the two experiments can be accounted for parsimoniously by Model_Log_ 2 that predicates response times on i. spatial attention, ii. general response probability and iii. spatially selective response probability. Critically, adding spatial signal probability per se as a predictor did not substantially increase the model evidence (Model_Log_ 3, Figure 3). These results suggest that spatial signal probability influences response times mainly indirectly via general and spatially selective response probabilities rather than as an independent factor.

Our findings indicate that the interactions between spatial attention and signal probability previously reported at the neural level^31^ are behaviourally relevant, yet they offer a new perspective: the interactions may represent ‘response related’ expectations encoding the general and spatially selective response probabilities. Observers concurrently form multiple sorts of expectations such as the expectation of a signal (i.e. signal probability), the expectation to make a response irrespective of (i.e. the general response probability) or dependent on (i.e. the spatial selective response probability) a particular signal location. Collectively, these expectations encode the probabilistic structure of sensory inputs and demands within the entire perception-action loop^37,38^. This sheds new light on the interplay of spatial attention and expectation. Current operational definitions of attention as task-relevance / response requirement and expectation as signal probability^1,35^ set up an artificial dichotomy between attention and signal probability. Yet signal probability and task-relevance are intimately linked by co-determining general and spatially selective response probabilities that shape response-related expectations and allocation of attentional resources. In the next step we therefore asked whether observers’ spatial allocation of attentional resources (i.e. main effect of attention on response times) is already accounted for by response-related expectations (i.e. general and spatially selective response probability). Using Bayesian model comparison we show that a more parsimonious model without spatial attention as an independent predictor performed substantially worse than the winning model (Model_Log_ 2). These results suggest that endogenous spatial attention can at least to some extent influence perceptual decisions above and beyond spatially-selective response probability. In line with this conjecture, observers have been shown to allocate attentional resources based on instructions alone^39,40^, even if response requirements are kept constant thereby dissociating attention and task-relevance. In order to dissociate additive and interactive effects of endogenous spatial attention (independent from task-relevance) and signal-related expectations at the neural level, future studies may manipulate endogenous spatial attention via instructions alone whilst holding the response requirements constant. Critically, our experimental results also reveal that the dichotomy between spatial expectation and spatial attention when operationally defined as response requirement over space may be useful from an initial analytical perspective. However, signal probability and attention as task-relevance are intimately linked by co-determining response-related expectations. Thus, changes in spatial expectation will inherently alter general or spatially selective response probabilities and therefore influence participant’s allocation of attentional resources over spatial hemifields.

From a cognitive perspective, changes in general response probability may be associated with increases in alertness, arousal or motor preparation^37,38^ leading to response facilitation. By contrast, spatially selective response probability needs to engage spatially selective mechanisms. For instance, the neural system may increase the precision of representations at spatial locations associated with a greater spatially selective response probability. Alternatively, observers may bias their decisional processes by shifting the starting point closer to the decisional boundary or lowering the decisional boundary of the evidence accumulation process^41,42^. In support of the latter, our signal detection theoretic analysis of response choices showed an increase in hits for higher general response probability in experiment 1 and a decrease in bias for higher spatially selective response probability in experiment 2. Critically, as participants always respond with the same effector organ (i.e. index finger of the dominant hand) to all visual and task-relevant auditory signals across both hemifields, decisional processes for all spatial locations need to finally map onto the same effector organ.

Cast in the predictive coding framework our results suggest that the brain iteratively adjusts its predictions of the sensory inputs at multiple levels across the cortical hierarchy. Critically, spatial attention as task-relevance profoundly modulates the precision of prediction errors and thereby the gain with which they impact higher cortical levels that are critical for response selection: First, the precision or gain of prediction errors is higher for sessions where many stimuli require a response (i.e. sessions with high general response probability). Second, the precision of prediction errors are selectively optimized for spatial locations associated with a high spatially selective response probability, i.e. locations where a high percentage of signals require a response. Guided by current task demands the brain thus adaptively optimizes the precision of spatial representations at locations that are critical for effective interactions.

Future neuroimaging and neurophysiological research will need to investigate whether the facilitatory effects of endogenous attention, signal probability, general and spatially selective response probability are mediated by similar or distinct neural mechanisms. For instance, they may be reflected in pre-stimulus baseline shifts or emerge only during post-stimulus processing (e.g. enhanced precision of spatial representations). Further, from a multisensory perspective we need to determine whether putative pre-stimulus baseline shifts, biases and increases in precision of spatial representations emerge only at higher levels of the cortical hierarchy associated with multisensory representations of space or in primary sensory areas across vision and audition either directly or via feed-back from higher order areas^43,44,45^.

In conclusion, our results across two experiments demonstrate that spatial attention and signal probability can influence signal detection either interactively or additively. These seemingly contradictory results can be reconciled by a new model that explains response times parsimoniously by spatial attention, general and spatially selective response probabilities. Our model provides a novel perspective on the intricate interplay of attention, signal- and response-related expectations in perceptual decisions, which is critical for effective interactions with the environment.

## General Methods

### Apparatus

During the experiment, participants rested their chin on a chinrest with the height held constant across all the participants. Auditory stimuli were presented at approximately 72 dB SPL, via HD 280 PRO headphones (Sennheiser, Germany). Visual stimuli were displayed on a gamma-corrected LCD monitor (2560 × 1600 resolution, 60 Hz refresh rate, 30” Dell UltraSharp U3014, USA), at a viewing distance of approximately 50 cm from the participant’s eyes. Stimuli were presented using Psychtoolbox version 3^46,47^ (www.psychtoolbox.org), running under Matlab R2014a (Mathworks Inc., Natick, MA, USA) on a Windows machine. Participants responded to all stimuli with the same index finger of their reported dominant hand. Responses were recorded via one key of a small keypad (Targus, USA). Throughout the experiment, participants’ eye-movements and fixations were monitored using Tobii Eyex eyetracking system (Tobii, Sweden).

### Stimuli

Spatial auditory stimuli (100 ms duration) were created by convolving a burst of white noise (with 5 ms onset and offset ramps) with spatially selective head-related transfer functions (HRTFs) based on the KEMAR dummy head of the MIT Media Lab^48^ (http://sound.media.mit.edu/resources/KEMAR.html).

Visual stimuli (i.e. the so-called ‘flash’) were white discs (radius: 0.88° visual angle, luminance: 196 cd/m2, 100 ms duration) presented on a grey background.

Both auditory and visual stimuli were presented at ±10° of visual angle along the azimuth (0° of vertical visual angle). A fixation cross was presented in the centre of the screen throughout the entire experiment.

In Experiment 2, two types of stimuli were presented, *target* and *non-targets*. Targets were identical to the non-targets, except that a brief gap (10 ms) was inserted after 45 ms.

Prior to the main experiment, participants were tested for their ability to discriminate left and right auditory stimuli in a brief series of 20 trials in a 2-alternative forced choice task. Participants indicated their spatial discrimination response (i.e. ‘left’ vs. ‘right’) via a two choice key press (group mean accuracy was 99.3% ± 1.7% SD in Experiment 1 and 98% ± 2.2% SD in Experiment 2).

## Experiment 1

### Participants

Sixteen healthy subjects (5 males, 11 females; mean age = 20.06 years; range 18-27 years; 15 right-handed) participated in experiment 1. The sample size was determined based on previous studies investigating attention/expectation^2-4,6,8-10,31,32^and/or multisensory integration^27,28^. All participants had normal or corrected to normal vision and reported normal hearing. In experiment 1, one participant was excluded post-hoc from the analysis because the overall performance accuracy on the target detection task was lower than 2 SD of the group mean (i.e. the across subjects’ mean = 98.2% correct). All participants provided written informed consent and were naïve to the aim of the experiment. The study was approved by the local ethics committee of the University of Birmingham (Science, Technology, Mathematics and Engineering (STEM) Ethical Review Committee) and the experiment was conducted in accordance with these guidelines and regulations.

### Design and procedure

Experiment 1 investigated the effect of auditory spatial attention and expectation on detection of auditory and visual targets using a 2 (*auditory spatial attention*: left vs. right hemifield) × 2 (*auditory spatial expectation*: left vs. right hemifield) × 2 (*stimulus modality*: auditory vs. visual) × 2 (*stimulus location*: left vs. right hemifield) factorial design (see Figure 1a). For the design figure, analysis and results, we pooled over the factor ‘stimulus location’ to provide a more succinct 2 (attention: *attended vs. unattended* hemifield) × 2 (expectation: *expected vs. unexpected hemifield*) × 2 (*stimulus modality*: auditory vs. visual) design. Auditory spatial expectation was manipulated as auditory spatial signal probability in the left and right hemifields across experimental sessions that were performed on different days. Observers were not informed about those probabilities, but learnt them implicitly. Auditory spatial attention was manipulated as ‘task-relevance’, i.e. the requirement to respond to an auditory target in the left vs. right hemifield over runs of 80 trials. Prior to each run a cue (duration: 2000 ms) informed the observer whether to respond to auditory targets either in the left or right hemifield. Critically, spatial attention and expectation were manipulated only in audition but not vision. Hence, observers needed to respond to all visual targets that were presented with equal probabilities in either spatial hemifields.

Each trial (SOA: 2300 ms) included three time windows (see Figure 1c): i. fixation cross alone (700 ms duration), ii. brief flash or sound (stimulus duration: 100 ms) and iii. fixation cross alone (1500 ms as response interval). Observers responded to the auditory stimuli in the attended hemifield and to the visual stimuli via key press with the same index finger (i.e. the same response for all auditory and visual stimuli) as fast and accurately as possible. They fixated the cross in the centre of the screen that was presented throughout the entire experiment with their fixation performance monitored via eye tracking.

Two sessions (i.e. spatial expectation left vs. right on different days) included 12 attention runs with 80 trials each. Runs were of two types: in run type A (coded in blue in Figure 1a and 1b) spatial attention and expectation were congruent (i.e. spatial attention was directed to the hemifield with higher auditory target frequency); in run type B (coded in green) spatial attention and expectation were incongruent (i.e. attention was directed to the hemifield with less frequent auditory stimuli). Thus, in total the experiment included 80 trials × 12 attention runs (6 runs of type A and 6 runs of type B) × 2 expectation sessions = 1920 trials in total. Specifically, each run type overall included 384 auditory non-targets for the expected hemifield (pooled over left and right) and 96 auditory non-targets for the unexpected hemifield (pooled over left and right). Each run type also included 240 visual non-targets for the expected hemifield and 240 visual non-targets for the unexpected hemifield (pooled over locations). For further details see Figure 1b which shows the absolute number of trials for each condition and run type.

The order of expectation (i.e. left vs. right) sessions was counterbalanced across participants; the order of attention runs was counterbalanced within and across participants and the order of stimulus location and stimulus modality were pseudo-randomized within each participant. Brief breaks were included after every two attention runs to provide feedback to participants about their performance accuracy (averaged across all conditions) in the target detection task and about their eye-movements (i.e. fixation maintenance).

Prior to each session, participants were familiarized with the stimuli in brief practice runs (with equal spatial signal probability) and trained on target detection performance and fixation (i.e. a warning signal was shown when the disparity between the central fixation cross and the eye-data samples exceeded 2.5 degrees). After the final session participants indicated in a questionnaire whether they thought the sound or the flash were presented more frequently in one of the two spatial hemifields. Fifteen out of the total 16 participants correctly reported that the auditory stimuli were more frequent in one hemifield and all participants reported the visual stimuli to be equally frequent across the two hemifields, suggesting that most participants were aware of the manipulation of signal probability in experiment 1.

## Experiment 2

### Participants

Twenty-four new healthy subjects (5 males, 19 females; mean age = 20.54; range 18-40 years; 20 right-handed) took part in experiment 2. The sample size was increased compared to experiment 1, because the number of targets (i.e. trials requiring a response) in the condition with the minimal number of trials was half the number of trials in the same condition of experiment 1 (48 versus 96, compare Figure 1b and 2b). All participants had normal or corrected to normal vision and reported normal hearing. Three participants were excluded post-hoc from the analysis because their overall performance accuracy on the target detection task was lower than 2 SD of the group mean (i.e. the across subjects’ mean = 98% correct). All participants provided written informed consent and were naïve to the aim of the experiment. The study was approved by the local ethics committee of the University of Birmingham (Science, Technology, Mathematics and Engineering (STEM) Ethical Review Committee) and the experiment was conducted in accordance with these guidelines and regulations.

### Design and procedure

Experiment 2 again investigated the effect of auditory spatial attention and expectation on detection of auditory and visual targets using a 2 (*auditory spatial attention*: left vs. right hemifield) × 2 (*auditory spatial expectation*: left vs. right hemifield) × 2 (*stimulus modality*: auditory vs. visual) × 2 (*stimulus location*: left vs. right hemifield) × 2 (*stimulus type*: target vs. non-target) factorial design (see Figure 2a). For the design figure, analysis and results, we pooled over the factor ‘stimulus location’ to provide a more succinct 2 (attention: *attended vs. unattended* hemifield) × 2 (expectation: *expected vs. unexpected hemifield*) × 2 (*stimulus modality*: auditory vs. visual) × 2 (*stimulus type*: target vs. non-target) design.

As in experiment 1, we manipulated spatial attention and expectation in audition alone and investigated its effect on response times to auditory and visual stimuli. Critically, in contrast to experiment 1, experiment 2 manipulated spatial expectation not directly via the frequency of auditory targets, but indirectly via the frequency of auditory non-targets across the two hemifields on different days. Thus, experiment 2 included two sets of stimuli for both auditory and visual modalities: targets and non-targets. The non-targets were identical to the stimuli in experiment 1. The targets were flashes or sounds with a 10 ms gap introduced after 45 ms, but otherwise identical to the non-targets. The non-targets never required a response in any sensory modality irrespective of hemifield. They were introduced to manipulate auditory stimulus expectation, i.e. auditory stimulus probability across the two hemifields, such that the general response probability was held constant across all conditions^32^ (see Figure 2a and below for further explanation). Again observers were not informed about those probabilities, but learnt them implicitly. Auditory spatial attention was manipulated as ‘task-relevance’, the requirement to respond to auditory targets in the left vs. right hemifield over runs of 96 trials. Prior to each run a cue (duration: 2000 ms) informed observers whether to respond to auditory targets either in the left or right hemifield. Critically, spatial attention and expectation were manipulated only in audition but not vision. Hence, observers needed to respond to all visual targets stimuli that were presented with equal probabilities in both spatial hemifields.

As shown in Figure 2c, the sequence and timing of a trial were identical to experiment 1. In experiment 2, observers responded only to the auditory targets (i.e. double sound) in the attended hemifield and the visual targets (i.e. double flash) via key press with the same index finger (i.e. the same response for all auditory and visual targets) as fast and accurately as possible.

As in experiment 1, two sessions (i.e. spatial expectation left vs. right on different days) included 12 attention runs with 96 trials each. Runs were of two types: in A (coded in blue in Figure 2a and 2b) spatial attention and expectation were congruent (i.e. spatial attention was directed to the hemifield with higher auditory target frequency), whereas in run B (coded in green) spatial attention and expectation were incongruent (i.e. attention was directed to the hemifield with less frequent auditory stimuli). Thus, in total the experiment included 96 trials × 12 attention runs (6 run of type A and 6 runs of type B) × 2 expectation sessions = 2304 trials in total. Specifically, each run type overall included 384 auditory non-targets for the expected hemifield (pooled over left and right) and 96 auditory non-targets for the unexpected hemifield (pooled over left and right). Each run type also included 240 visual non-targets for the expected hemifield and 240 visual non-targets for the unexpected hemifield (pooled over locations). Further, each run type included 48 auditory and 48 visual targets for the expected hemifield and 48 auditory and 48 visual targets for the unexpected hemifield. For further details see Figure 2b which shows the absolute number of trials for each condition and run.

The procedural details (e.g. counterbalancing, eye tracking, post-questionnaire) were otherwise comparable to experiment 1. Eight out of 24 participants correctly reported that the auditory stimuli were more frequent in one hemifield and 23 out of 24 participants reported the visual stimuli to be equally frequent across the two hemifields, suggesting that most participants were not explicitly aware of the manipulation of signal probability in experiment 2. Most likely, participants were less aware of the expectation manipulation in experiment 2 than experiment 1, because signal probability was manipulated only for the task-irrelevant non-targets. Further, the inclusion of target and non-target stimuli in the same paradigm increased the stimulus variability making the expectation manipulation less apparent.

### Experiments 1 and 2: Spatial signal, general response and spatially selective response probability

Both experiments 1 and 2 orthogonally manipulated spatial attention (i.e. response requirement) and expectation (i.e. spatial signal probability). Yet, while experiment 1 manipulated spatial signal probability (i.e. the probability of a signal irrespective of sensory modality in the left or right hemifield) directly via the probability of task-relevant auditory targets across hemifields (see Figure 1d, left bar plot and Supplementary Table S1), experiment 2 manipulated it indirectly via additional task-irrelevant auditory non-targets that never required a response (see Figure 2d, left bar plot and Supplementary Table S1). As a result, the two experiments were associated with different profiles of i. the general response probability (i.e. the probability that the observer needs to make a response irrespective of the hemifield in which the signal is presented, see Figure 1d and 2d, middle bar plot and Supplementary Table S1) and ii. the spatially selective response probability (i.e. the probability that the observer needs to make a response conditioned on that the signal is presented in a particular hemifield, see Figure 1d and 2d, right bar plot and Supplementary Table S1).

In experiment 1, the general response probability is greater in the run type A (coded in blue in Figure 1d, middle bar plot), where spatial attention and expectation are congruent (i.e. spatial attention is directed to the hemifield with higher auditory target frequency), than in run type B (coded in green), where spatial attention and expectation are incongruent (i.e. attention is directed to the hemifield with less frequent auditory stimuli). Likewise, the spatially selective response probability changes across conditions co-determined by auditory spatial attention and spatial signal probability (Figure 1d, right bar plot).

Experiment 2 manipulated auditory spatial expectation via additional non-targets that never require a response. As a consequence, the general response probability is held constant throughout the entire experiment (see Figure 2d, middle bar plot). Further, the increase in spatial signal probability in the expected hemifield inherently decreases the spatially selective response probability (see Figure 2d, right bar plot).

## Experiment 1 and 2: Data analysis

### Eye movement analysis

We excluded trials where participants did not successfully fixate the central cross based on a dispersion criterion (i.e. distance of fixation from subject’s centre of fixation as defined in calibration trials > 1.3 degrees for three subsequent samples^49^) (an additional analysis including all trials yielded basically equivalent results). Our eye tracking data confirmed that participants successfully maintained fixation in both experiments with only a small number of trials to be excluded (experiment 1: excluded auditory response trials 2.9% ± 1% [across subjects mean ± SEM]; excluded visual response trials 2.8% ± 1% [across subjects mean ± SEM]; experiment 2: excluded auditory response trials 4.4% ± 1.6% [across subjects mean ± SEM]; excluded visual trials 4.6% ± 1.5% [across subjects mean ± SEM].

### Response choice and time analysis

Using signal detection theory we reported hit rates for experiment 1 and experiment 2 and d-prime (Z(hit rate) - Z(false alarm rate)) as well as bias (−0.5[Z(false alarm rate) + Z(hit rate)]) for the visual modality for experiment 2. D-prime and bias could not be computed for experiment 1 (because non-target trials were not included in the paradigm) and for auditory modality of experiment 2 (because participants did not make any false alarms in the +att-exp condition). Response time analysis was limited to correct trials only and response times within the range of participant- and condition-specific mean ± two SD and <1500 ms.

#### General linear models fitted independently for each experiment

Inference was initially made separately for each experiment at the random effects level to allow for generalization to the population. For auditory targets in the attended hemifield, condition-specific hit rate (experiment 1), d-prime and bias (experiment 2), and median response times (experiment 1 and 2) for each subject were entered into a two-sided paired t-tests with *expectation* (expected, unexpected) as factor. For visual targets, condition-specific hit rate (experiment 1), d-prime and bias (experiment 2), and median response times (experiment 1 and 2) for each subject were entered into a 2 (*attention*: attended, unattended) × 2 (*expectation*: expected, unexpected) repeated measures analysis of variance (ANOVA). Please note that the computation of bias and d-prime was not possible for experiment 1, because participants had to respond to all visual stimuli. Unless otherwise indicated, we only report effects that are significant at *p* < 0.05.

#### Generalized mixed effects model fitted jointly to both experiments

To identify a model that parsimoniously explains the data jointly from both experiments, we used generalized mixed effects models across both experiments and Bayesian model comparison. We specified six generalized mixed effects models that were organized in a 3 (combination of *predictors*, as in Figure 1d and 2d and Supplementary Table S1) × 2 (*probabilities*: linear vs. log-transformed) factorial model space. The first factor specified the set of fixed effects predictors:

Traditional Model 1: i. spatial attention: categorical dummy variable encoding whether or not stimulus is presented in the attended hemifield, ii. spatial expectation: signal probability of the hemifield where the stimulus is presented, iii. interaction between spatial attention and expectation.

Model 2: i. spatial attention: categorical dummy variable encoding whether or not stimulus is presented in attended hemifield, ii. general response probability: probability that the participants need to respond to the stimulus prior to knowing where the stimulus is presented, i.e. this probability does not depend on the spatial location of the stimulus, iii. spatially selective response probability: probability that the participants need to respond given that the stimulus is presented in a particular hemifield, i.e. this probability depends spatial location.

Model 3: Model 2 with the added regressor iv. spatial expectation: signal probability of the hemifield where the stimulus was presented.

The second factor specified whether the probabilities encoded in the regressor predicted response times in a linear or log-transformed fashion to accommodate previous evidence that response times may be non-linearly related to stimulus probability^34^.

All six models also included a fixed effects regressor encoding the sensory modality (i.e. auditory vs. visual) of the stimulus. Furthermore, they modelled subject and experiment as random effects to account for the higher order organization into two experiments.

We fitted these six models jointly to the data from experiment 1 and 2 using maximum likelihood estimation and compared the non-nested models using the Bayesian information criterion (BIC) as an approximation to the model evidence. We fitted the winning model again to the data from both experiments using restricted maximum likelihood estimation to obtain unbiased estimates for generating the model predictions shown as grey crosses in Figure 1e and 2e (see guidelines in^50^).

## Acknowledgements

This research was funded by ERC-2012-StG_20111109 multsens.

## Author Contributions

A.Z. and U.N. conceived and designed the experiments. A.Z. performed the experiments. A.Z. and U.N. analysed the data. A.Z. and U.N. contributed to the writing of the manuscript.

## Competing financial interests

The authors declare no competing interests.

