## Supplementary Materials for "Additive and interactive effects of spatial attention and expectation on perceptual decisions"

### Supplementary information

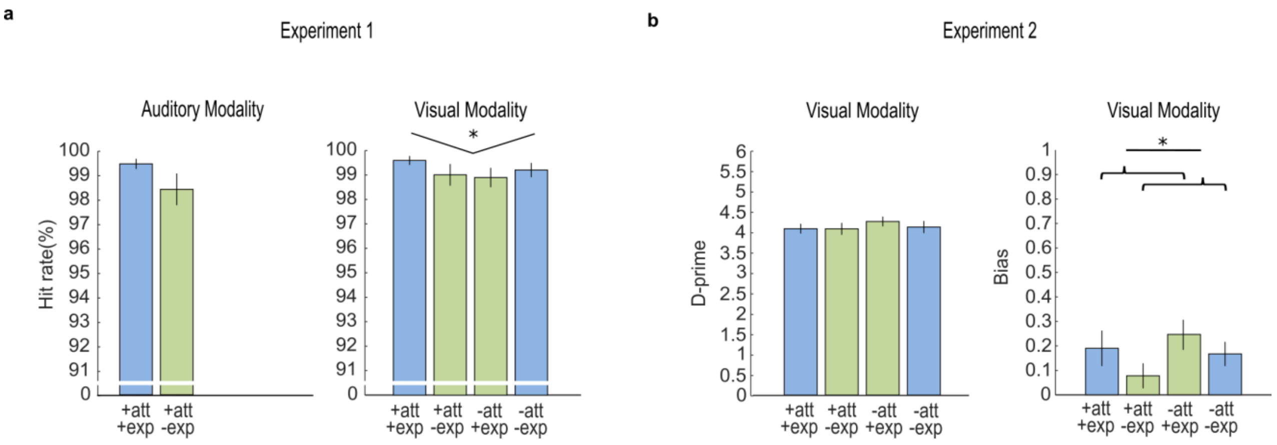


**Figure S1 | Experiment 1: hit rates; Experiment 2: d-prime and bias**

**a.** Experiment 1: hit rates for each of the six conditions with response requirements. The brackets and stars indicate significance of main effects and interactions. * $p < 0.05$.

**b.** Experiment 2: d-prime and bias values for each of the four conditions for the visual modality. The brackets and stars indicate significance of main effects and interactions. * $p < 0.05$.

**Table S1.** Spatial signal probability, general response probability and spatially selective response probability for each condition in experiment 1 and experiment 2.

S = signal location, $H_{..att..exp}$ = hemifield e.g. +att +exp, R = response.

| | Conditions | | | |
|---|---|---|---|---|
| | +att +exp | +att -exp | -att +exp | -att -exp |
| **Experiment 1** | | | | |
| Spatial signal probability $P(S = H_{..att..exp})$ | 0.65 | 0.35 | 0.65 | 0.35 |
| General response probability $P(R = +)$ | 0.9 | 0.6 | 0.6 | 0.9 |
| Spatially selective response probability $P(R = + \mid S = H_{..att..exp})$ | 1 | 1 | 0.384 | 0.714 |
| **Experiment 2** | | | | |
| Spatial signal probability $P(S = H_{..att..exp})$ | 0.625 | 0.375 | 0.625 | 0.375 |
| General response probability $P(R = +)$ | 0.123 | 0.123 | 0.123 | 0.123 |
| Spatially selective response probability $P(R = + \mid S = H_{..att..exp})$ | 0.131 | 0.218 | 0.065 | 0.109 |

Please note that these probabilities are computed pooled over auditory and visual modalities.

The spatial signal probability is the probability that a signal is presented in a particular hemifield, e.g. in experiment 1: P = 0.65 for the hemifield that is attended and expected vs. P = 0.35 for the hemifield that is unattended and unexpected (n.b. the spatial signal probabilities sum to one for the two conditions presented in the same run).

The general response probability is the probability that a trial requires a response in a particular run irrespective of where the signal is presented, e.g. in experiment 1: P = 0.9 for runs where attention and expectation are congruent vs. P = 0.6 for runs where expectation and attention are incongruent.

The spatially selective response probability is the probability that a response is required conditioned on the signal being presented in a particular hemifield, e.g. in experiment 1: the probability of a response is P = 0.384, if the signal is presented in a hemifield that is not attended, but expected.

As a result, we can compute the general response probability by multiplying the spatially selective response probability with the spatial signal probability and marginalizing over the spatial signal probability; e.g.

$$P(R = +) = P(R = + | S = H_{+att+exp})\, P(S = H_{+att+exp}) + P(R = + | S = H_{-att-exp})\, P(S = H_{-att-exp})$$
